# Dynamic architecture of the *Escherichia coli* Structural Maintenance of Chromosomes (SMC) complex, MukBEF

**DOI:** 10.1101/547786

**Authors:** Karthik V. Rajasekar, Minzhe Tang, Rachel Baker, Katarzyna Zawadzka, Oliwia Koczy, Florence Wagner, Jani Reddy Bolla, Carol V. Robinson, Lidia K. Arciszewska, David J. Sherratt

## Abstract

Structural Maintenance of Chromosomes (SMC) complexes use a proteinaceous ring-shaped architecture to organise chromosomes, thereby facilitating chromosome segregation. They utilise cycles of ATP binding and hydrolysis to transport themselves rapidly with respect to DNA, a process requiring protein conformational changes and multiple DNA contacts. We have analysed changes in the architecture of the *Escherichia coli* SMC complex, MukBEF, as a function of nucleotide binding to MukB and subsequent ATP hydrolysis. This builds upon previous work showing that MukF kleisin directs formation of a MukBEF tripartite ring as a consequence of functional interactions between the C- and N-terminal domains of MukF with the MukB head and neck, respectively (Zawadzka et al., 2018). Using both model truncated substrates and complexes containing full length MukB, we now demonstrate formation of MukBEF ‘dimers of dimers’, dependent on MukF dimerization, MukB head-engagement and MukE, which plays an essential role in organizing MukBEF complexes.

## Introduction

Structural Maintenance of Chromosomes (SMC) complexes, which are present in all domains of life, share a distinctive architecture in which a tripartite proteinaceous ring is formed by a dimer of two SMC molecules and a kleisin that connects the two SMC ATPase heads. Interactions of a kleisin C-terminal domain with the cap of an SMC head and the kleisin N-terminal region with a coiled-coiled ‘neck’ adjacent to the head of the partner SMC molecule lead to this connection (Figure 1; Bürmann et al., 2013, Gligoris et al., 2014; Huis in ‘t Veld et al., 2014; Zawadzka et al., 2018). Emerging evidence supports the view that SMC complexes are mechanochemical motors that use cycles of ATP binding and hydrolysis to transport themselves rapidly with respect to DNA, extruding DNA loops during this transport (Ganji et al., 2018). Such activities have important roles in chromosome organization-individualisation and segregation, as well as other aspects of DNA management (Nolivos and Sherratt, 2014; Hirano, 2016; Uhlmann, 2016; Nasmyth 2017).

**Figure 1.**
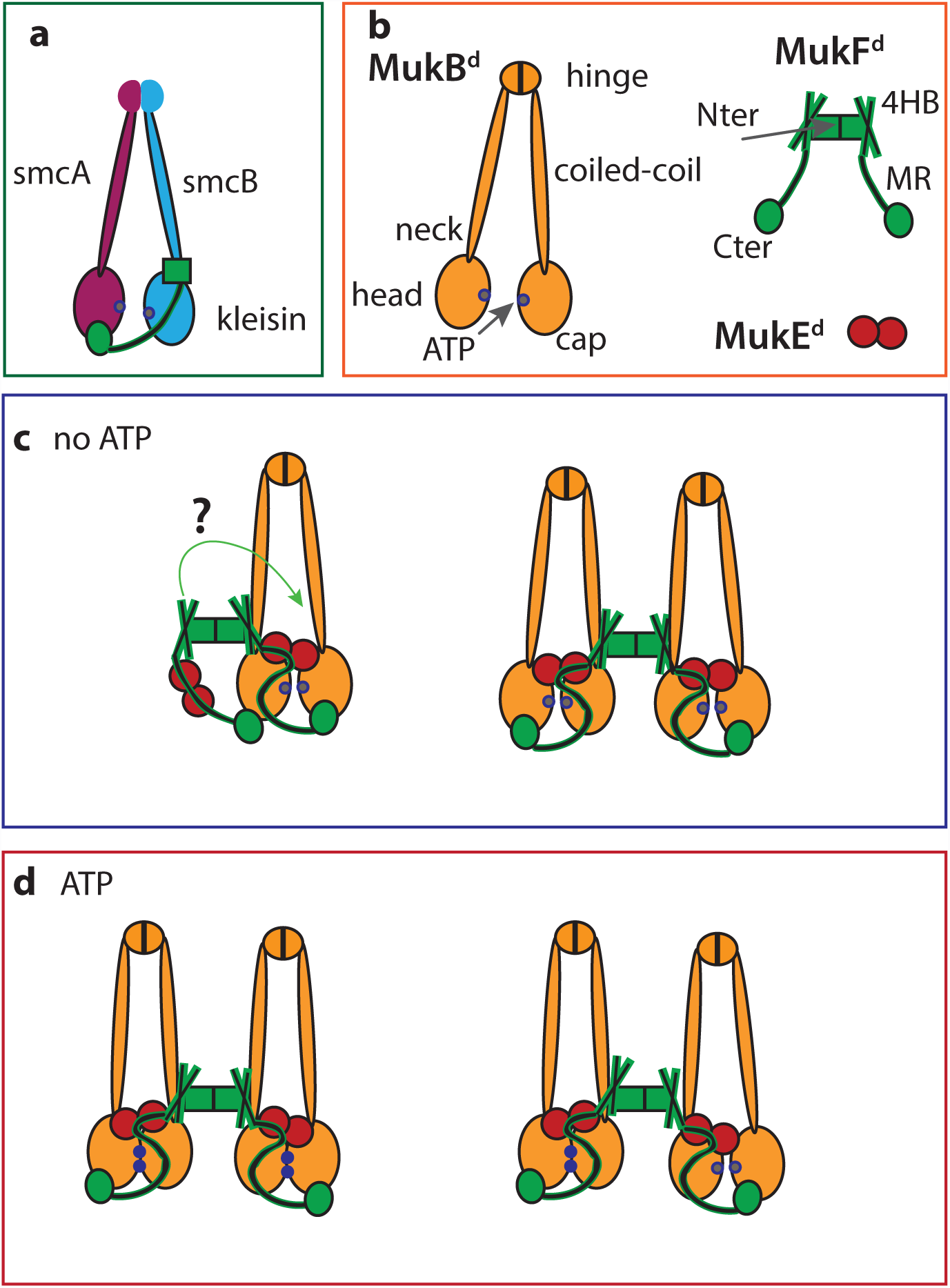
Schematic showing conserved SMC architectures and possible MukBEF stoichiometries and architectures. Panel **a**, generic SMC complex architecture showing the tripartite proteinaceous ring formed by the kleisin and SMC proteins. **b**, the MukBEF complex. **c**, **d** possible and proposed MukBEF architectures, without (**c**) and with (**d**) ATP. In **d**, the right panel shows a possible complex after ATP is hydrolysed in one of the dimers of a dimer of dimer complex. When heads engage in the presence of ATP, the MukF middle region blocks the binding of a second MukF C-terminal domain to the second MukB head within a MukB dimer (Woo et al., 2009). When heads are unengaged, each MukB head can bind a MukF C-terminal domain, potentially leading to ‘daisy chain’ forms of a higher complexity than dimers and dimers of dimers (not shown). ATP-bound ATPase active sites denoted as blue dots on the heads and ADP-bound or nucleotide-unbound as grey dots.

Although the *Escherichia coli* SMC complex, MukBEF, shares many aspects of the distinctive SMC complex architecture, its kleisin, MukF, is dimeric, which could potentially facilitate the formation and action of higher order complexes (Fennell-Fezzie et al., 2005; Badrinaryananan et al., 2012; Nolivos and Sherratt, 2014). MukBEF homologs are only found in a fraction of γ-proteobacteria, where they have co-evolved with a group of other proteins, including MatP, Dam and SeqA (Brézellec et al., 2006). MukBEF also coordinates the localization and action of TopoIV (Nicolas et al., 2014; Zawadzki et al., 2015), with MatP-*matS* regulating the distribution and activity of both MukBEF and TopoIV in cells (Nolivos et al., 2016).

In E*.coli* cells, ~200 dimeric MukBEF complexes (or their multimeric equivalent) are present, with ~ 40% of these being tightly associated with chromosomal DNA, of which 30-50% form clusters in which the functional units are dimers of MukBEF dimers or multiples thereof (Badrinarayanan et al., 2012). Clusters of wild-type MukBEF complexes are positioned at mid-cell in new born cells and the cell quarter positions thereafter by a ‘phase-locked Turing pattern’ (Murray and Sourjik, 2017). These clusters position the chromosome replication origin region (*ori*) (Danilova et al., 2007; Nolivos et al., 2016; Hoffman et al., 2018), thereby facilitating chromosome organisation and segregation. ATP binding and MukB head engagement are essential for the formation of MukBEF clusters, as they are present in wild type and in hydrolysis-deficient mutants (MukB^EQ^) cells, but not in cells impaired in nucleotide binding or in head engagement (Badrinarayanan et al., 2012).

To help understand how MukBEF performs its functions in chromosome management, we analysed changes in the architecture and stoichiometry of MukBEF complexes *in vitro* as a function of nucleotide binding and hydrolysis. Using a combination of biochemical and biophysical approaches on truncated and then full-length MukBEF complexes, we have demonstrated that dimers of head-engaged MukBEF dimers form *in vitro* when bound to AMPPNP, a non-hydrolysable analogue of ATP, or to ATP when hydrolysis is impaired. We have shown the role of MukE in formation of these dimers of dimers and present insight into the architectures of complexes with engaged and unengaged heads.

## Results

### MukF dimers direct formation of dimers of heads-engaged MukB dimers

To reveal the architectures and stoichiometries of MukBEF complexes experimentally, a truncated derivative of MukB, MukB_HN_, (MukB Head-Neck, subsequently abbreviated as HN) containing the MukB ATPase head and ~30% of the adjacent coiled-coil, was used in initial biochemical analyses. This coiled-coil contains the ‘neck’ to which a MukF 4-helix bundle, adjacent to the N-terminal dimerization domain, binds and activates MukB ATPase (Figure 1; Zawadzka et al., 2018). This strategy was chosen initially because of the technical challenges of incisive *in vitro* analysis of large ~1 MDa full length MukBEF complexes. A MukF dimer has four independent interfaces for binding MukB; the two MukF C-terminal domains and two N-terminal 4-helix bundles, which bind the MukB head and neck respectively (Figure 1). Therefore, each MukF dimer could bind from two to four MukB molecules.

HN formed complexes with MukEF, in the presence of AMPPNP, a non-hydrolysable analog of ATP, in size exclusion chromatography-multiangle light scattering (SEC-MALS) (Figure 2A). The broad red peak was predicted to be composed of two major components; material in the leading edge having a mass of 550 kDa (red square) and material in the lagging edge (red spot) with a mass of ~404 kDa. Complexes of these masses correspond to a 4HN-2F-4E complex (red square) and a 2HN-2F-4E-complex (red spot), respectively. The former complex is equivalent to a dimer of dimers MukBEF complex when MukB is a full-length wild-type dimer (Figure 1). SEC-MALS of samples with ADP revealed just the presence of the ~407 kDa complex, the mass of a 2HN-2F-4E complex (blue spot), which is equivalent to a dimeric MukBEF complex. Consistent with this interpretation, native gel electrophoresis demonstrated the AMPPNP-dependent formation of a slower moving complex (Figure 2A; upper panel; red square), along with faster running putative 2HN-2F-4E complexes formed in the presence of ADP (blue and red spots). Therefore, both the SEC-MALS and native gels demonstrate the formation of putative dimer of engaged-head dimer complexes, dependent on AMPPNP. Incubation with ATP gave the same electrophoretic profile as ADP, presumably because the ATP in any given complex was hydrolysed before analysis under the conditions used. A HN^SR^ derivative that is deficient in head-engagement (Badrinarayanan et al., 2012; Lammens et al., 2004) failed to give the equivalent dimer of dimer complexes on addition of AMPPNP in both SEC-MALS and native electrophoresis (Figure 2B), thereby providing further support for the interpretation that MukB head engagement is required for the formation of dimer of dimer complexes. Analysis of HN^EQ^, which binds ATP but is impaired in hydrolysis as a consequence of the Walker B motif mutation (Badrinarayanan et al., 2012; Lammens et al., 2004), showed that it forms the equivalent of dimer of dimer complexes in the presence of MukEF and ATP (Figure 2C), supporting our interpretations. Control experiments showed that HN was monomeric because of the lack of a dimerization hinge, while MukF and MukE were dimeric, as expected (Figure 2D).

**Figure 2.**
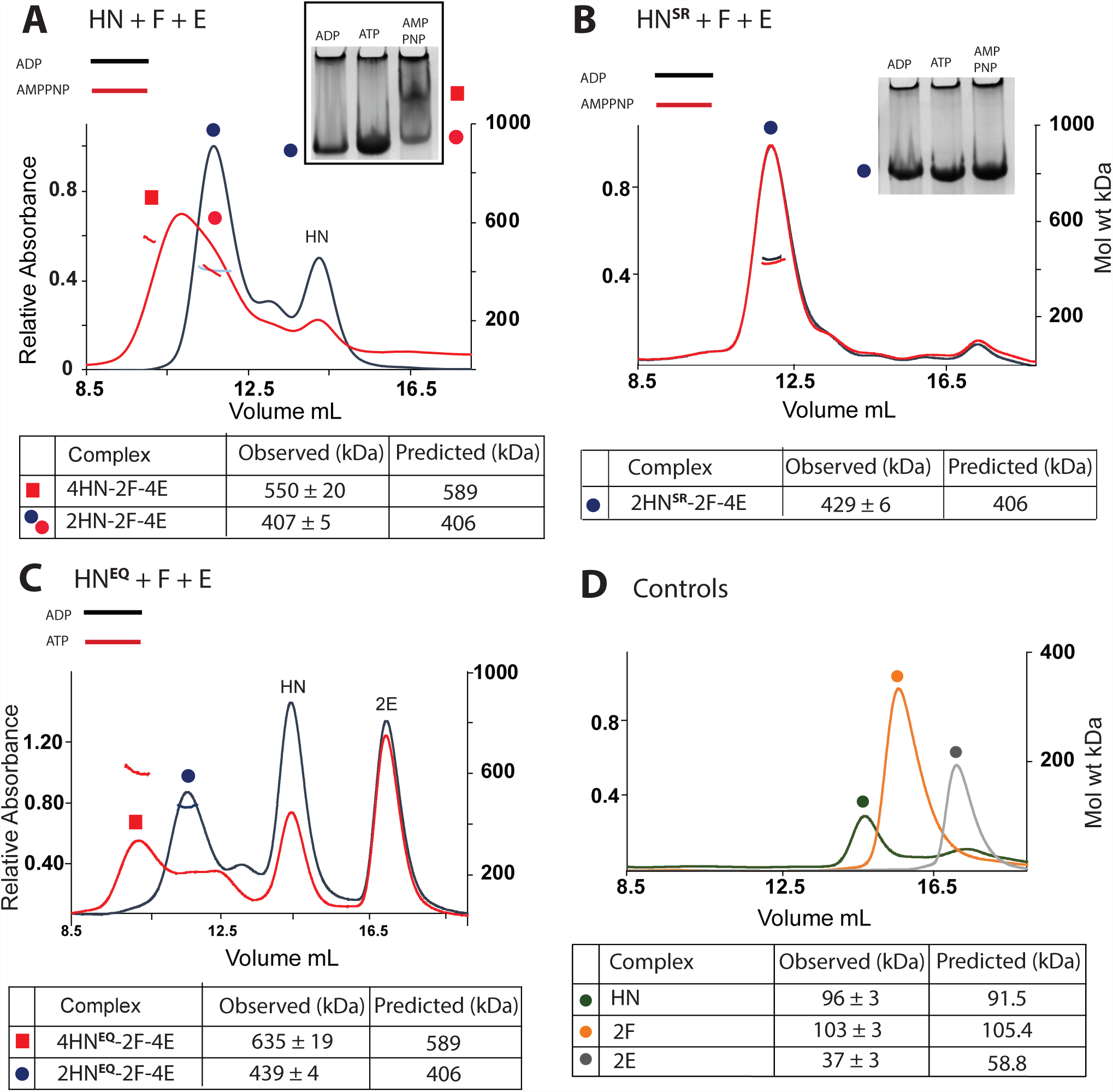
MukB head engagement is required for the formation of dimer of dimer MukBEF complexes. (**A-C**) Native PAGE (A, B) and SEC-MALS (A-C) analyses of the stoichiometry of HN/HN^SR^/HN^EQ^ complexes with MukFE in the presence and absence of ATP/AMPPNP. 10μM HN, 5μM F and 10μM E were incubated for 3 hours at room temperature with ADP or AMPPNP/ATP (1mM) prior to loading onto a 6% native gel, or a Superose 6 column. (**D**) SEC-MALS analysis of individual Muk proteins. Predicted and observed masses of the complexes are tabulated below with the values and their uncertainties derived from a single representative SEC-MALS experiment.

To test whether interactions of HN with both the MukF C-terminal domain and the 4-helix bundle adjacent to the MukF N-terminal domain are necessary to form dimer of dimer complexes, MukEF were incubated with HN^C*^, which is deficient in interaction with the MukF C-terminal domain (Figure 3A). Only a trace of 4HN^C*^-2F-4E complexes was observed in the presence of AMPPNP (filled red square), thereby demonstrating that interaction of the MukF C-terminal domain with the MukB cap is essential for formation of dimers of dimers. Supporting this conclusion, incubation of HN with FN10, lacking the MukF C-terminal domain, and MukE (Figure 3A; Zawadzka et al., 2018), gave no dimer of dimers complexes in the presence of AMPPNP. Interaction of MukEF with the MukB head (H), lacking the neck, or a mutant in the neck that impairs interaction with the MukF 4-helix bundle (HN^N*^; Zawadzka et al., 2018), also led to a much reduced level of dimer of dimer complexes in the presence of AMPPNP (unfilled and filled red squares, respectively). We conclude that interaction of both the MukF 4-helix bundle with the MukB neck and the interaction of the MukF C-terminal domain with the cap on the MukB head are crucial for efficient dimer of dimer formation, with the MukF C-terminal interaction with the head having a more important role than the interaction of the MukF 4-helix bundle with the neck.

**Figure 3.**
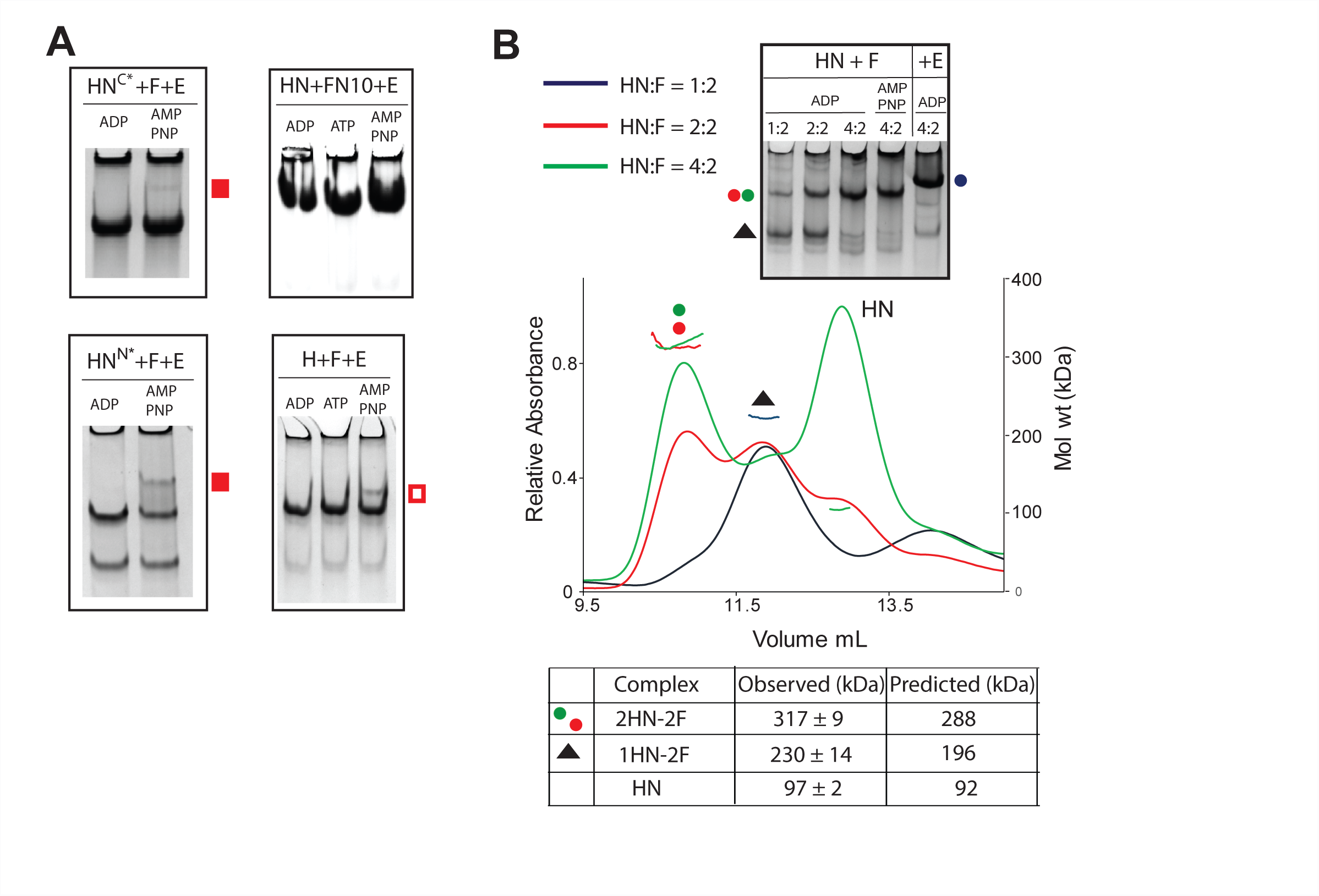
Dimer of dimer formation requires MukE, and HN interactions with both the MukF C-terminal domain and the MukF 4-helix bundle. (**A**) Native PAGE of complexes generated with HN and F variants deficient in binding across one (HN^C*^, FN10), or the other (HN^N*^and H), HN-F interface. The position of low levels of dimer of dimer complexes is indicated with filled red squares (HN), or an unfilled red square (H). HN^C*^ carries the following aa residues alterations: F1453S, H1458A, R1465A (**B**) SEC-MALS analyses of HN-F complexes at different HN:F ratios; 2.5/5/10 μM HN was mixed with 5μM F prior to separation through a Superdex 200 column. Predicted and observed masses of the complexes are tabulated below. The same mixtures incubated with ADP along with samples containing 10μM HN+5μM F and either AMPPNP or 10μM E were analyzed on 6% native gels.

We then addressed whether MukE is required to form AMPPNP-dependent heads-engaged dimer of dimer complexes and how lack of MukE influenced the stoichiometry of complexes. Incubation of HN with MukF dimers at varying molar ratios gave complexes having a molecular mass of ~317 kDa in the presence of ADP, close to the mass expected of 2HN-2F complexes (Figure 3B; red and green spots). At a ratio of HN: MukF of 0.5, most material ran with the mass predicted for HN-2F complexes (black triangle). In native gels, the same titrations in the presence of ADP showed the formation of a slower migrating complex, which increased in abundance as relative HN concentration increased; we interpret these complexes as 2HN-2F. Because replacement of ADP by AMPPNP made little difference to the complexes’ mobility, we conclude that MukE is required to form dimer of dimer complexes. The native gel also shows how MukE influences the mobility of HN-MukF complexes (compare blue spot with red/green spots).

To confirm that two MukE dimers bind tightly to a MukF dimer, we used SEC-MALS and native gels (Figure 3, figure supplement 1A), along with isothermal calorimetry (ITC) (Figure 3, figure supplement 1B). These analyses showed that dimeric MukF binds two MukE dimers and gave a K_d_ of 6.97 ± 2.6 nM for the interaction. We failed to detect any interaction of MukE with the MukB head in biochemical analyses (Figure 3, figure supplement 2), despite crystal structures showing interaction surfaces between MukE and MukB heads (Woo et al., 2009).

Although the assays described here, are suitable for detecting the equivalent of MukBEF dimers of dimers, as judged by dimeric MukF molecules capturing four molecules of HN, complexes containing a dimeric MukF molecule bound by two HN molecules could correspond to either dimeric full length MukBEF complexes, or to dimers of MukBEF dimers. Subsequent experiments were designed to help resolve this ambiguity.

### Further characterization of MukBEF architecture and stoichiometry

To further characterize MukBEF architecture, we analyzed the interaction of HN with two different MukF derivatives. Dimeric FN10, lacking the MukF C-terminal domain (Figure 4A, top), interacts normally with MukE (Zawadzka et al., 2018). Incubation with HN demonstrated complexes of mass expected for FN10 dimers with one or two bound HN molecules, (Figure 4A, yellow and grey spots). The equivalent complexes (yellow and grey spots) were inferred from native gel electrophoresis. The proportion of complexes with two HN molecules increased as the relative concentration of added HN increased in both SEC-MALS and native gel electrophoresis. We conclude that in the absence of MukE, that the two 4-helix bundles of a FN10 dimer can each bind one HN neck (Figure 4D; panel **a**, bottom).

**Figure 4.**
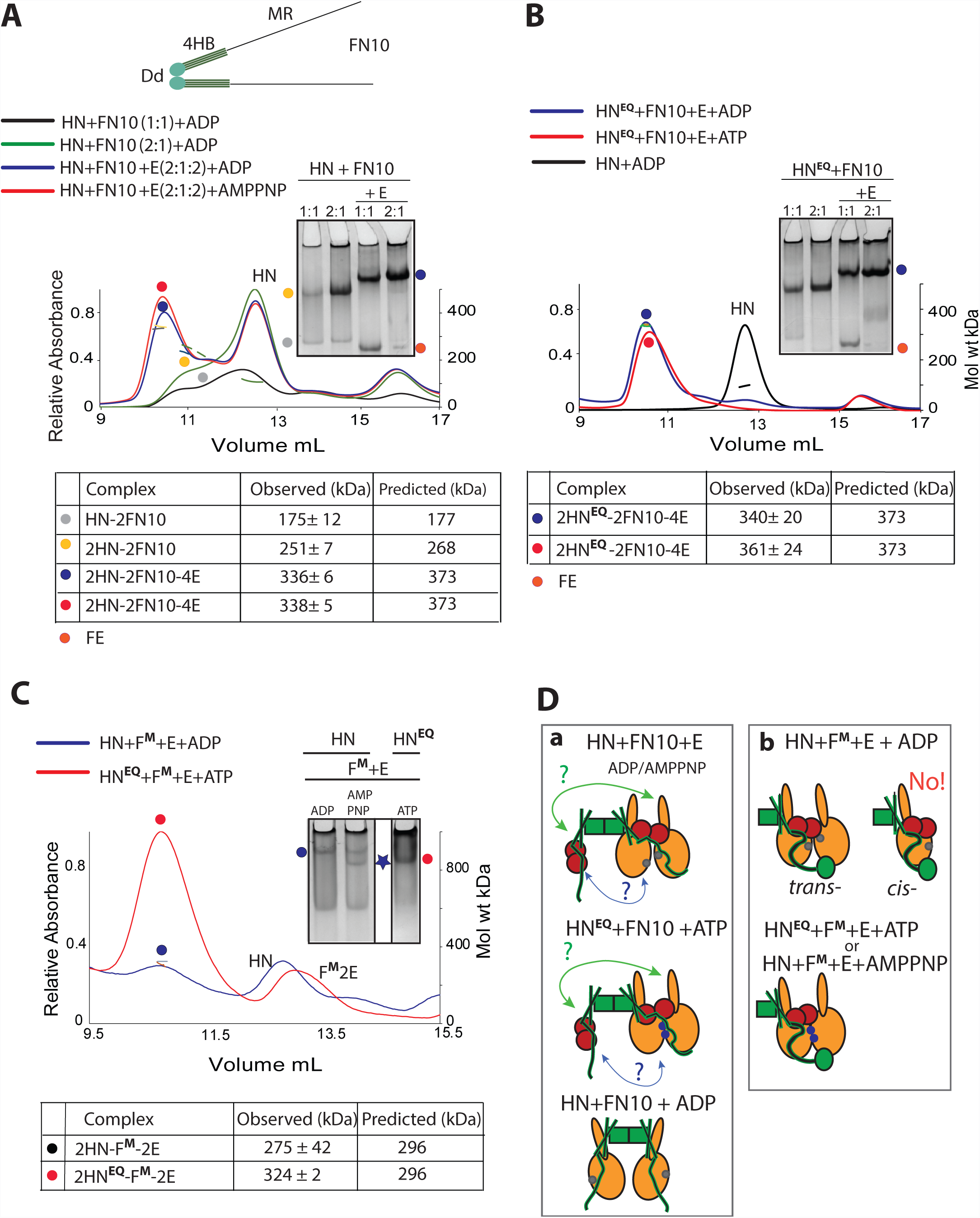
Architecture of MukBEF complexes. (**A, B**) Top, schematic of FN10 (Dd, N-terminal dimerization domain; 4HB, 4-helix bundle; MR, middle region; the C-terminal domain is deleted). Below, SEC-MALS and native PAGE analyses of HN and HN^EQ^ complexes generated with FN10 dimers. For SEC-MALS samples at the indicated protein ratios were incubated with ADP, AMPPNP or ATP before separation through a Superose 200 column. FN10 was at 5μM and E, when present, was 10μM. Predicted and observed masses of complexes are tabulated below. Native gel samples were incubated with ADP (**A**) or ATP (**B**) prior to loading on a gel. (**C**) SEC-MALS and native PAGE analyses of HN and HN^EQ^ complexes generated with FM-E in the presence of ADP, AMPPNP, or ATP as indicated. The proteins were at concentrations 10μM HN/HN^EQ^, 5μM F^M^ and 10μM E. (**D**) Schematics of the proposed architectures with FN10 (panel **a**) and MukF^M^ (panel **b**). The green arrows (**a**) indicate a possible interaction between the FN10 4HB and the neck of the distal HN molecule. A second potential interaction between the ‘free’ FN10 middle region and its bound MukE to the proximal HN molecule is indicated by blue arrows. The bottom cartoon in (**a**) shows two HN molecules binding a FN10 dimer through interactions with the 4HBs. ATP-bound ATPase active sites denoted as blue dots on the heads and ADP-bound or nucleotide-unbound as grey dots.

Although, the presence of MukE did not alter the relative stoichiometry of FN10 with HN, it led to a higher proportion of complexes with two HN molecules bound, indicating that MukE stabilises these complexes. The SEC-MALS profiles were the same in ADP and AMPPNP, consistent with the earlier conclusion that interactions of the MukF C-terminal domain with the HN cap are required for dimer of dimer formation. Equivalent SEC-MALS and native gel profiles were observed when HN^EQ^ in the presence of ATP was used rather than HN. (Figure 4B). Since HN^EQ^ is predominantly dimeric upon incubation with ATP (Figure 4, supplementary figure 1), we conclude that only one pair of heads-engaged HN molecules can bind to a FN10 dimer in the presence of MukE, either because the necks of the two HN molecules in these complexes occupy both 4-helix bundles (Figure 4D; panel **a**, green arrows), or because binding of a HN neck to one 4-helix bundle prevents the second 4-helix bundle in a dimer interacting with a another neck because of conformational changes/steric clashes. We do not know if a ‘free’ FN10 middle region with bound MukE can interact with the second HN molecule (**a;** blue arrows).

Then we used a mutated MukF derivative that is unable to form dimers because of amino acid substitutions in the dimerization domain (MukF^M^), although it has an intact 4-helix bundle, MukE binding site and C-terminal domain. MukF^M^ was incubated with either HN or HN^EQ^ and MukE (Figure 4C). SEC-MALS showed the formation of complexes containing two HN molecules bound to a single MukF monomer, with HN^EQ^-ATP giving a much higher fraction of such complexes (red spot), as compared to HN-ADP (blue spot) This result was corroborated by the native gel profiles. Therefore, in the presence of MukE, a single MukF monomer, with an intact C-terminal domain and 4-helix bundle, can bind two HN molecules, irrespective of whether they are in the heads engaged (HN^EQ^-ATP or HN-AMPPNP), or unengaged state (HN-ADP). Unsurprisingly, the native gel shows a higher proportion of such complexes is present when the heads are engaged, with the complexes having a higher mobility, indicative of a more compact conformation (blue star). This result argues that a single HN molecule is unable to engage both its neck and cap with the 4-helix bundle and C-terminal domain of the same MukF polypeptide in the presence of MukE, which likely plays a role in directing this arrangement. Otherwise MukF monomers bound by a single HN molecule would be the dominant species. Figure 4D summarizes the proposed architectures that are demonstrated when HN are complexed with FN10 in the absence and presence of MukE (panel **a**), or when HN was complexed with MukF monomer in the presence of MukE (panel **b**). The failure to observe the ‘*cis*-configuration’ in which the neck and head of a single MukB molecule bind both the 4-helix bundle and C-terminal domain of a single MukF monomer (panel **b**) is consistent with the *trans*-configuration being important in directing a tripartite proteinaceous SMC ring (Figure 1). The observation that a single heads-engaged HN^EQ^ dimer binds to both a FN10 dimer and a MukF monomer (Figure 4D; compare panels **a** and **b**) is consistent with the conclusion above that FN10 dimers bound by MukE can only bind two HN molecules irrespective of whether they are engaged or not (**a**).

### Full length MukB forms dimers of heads-engaged dimer complexes with MukEF and AMPPNP

To ascertain whether the AMPPNP- and MukE-dependent formation of the equivalent of dimer of dimer complexes, characterized with truncated MukB, could be observed for complexes made with intact MukB, we analyzed full-length MukBEF complex by SEC-MALS (Figure 5A). These MukBEF complexes were not resolvable by native gel electrophoresis. Furthermore, available size exclusion columns had limited resolution in the size range expected for full-length MukBEF complexes (0.5-1 MDa). Nevertheless, a Superose 6 column, combined with MALS, gave sufficient resolution to observe changes in mass and conformation, although mass predictions from SEC-MALS over-estimated the theoretical masses by ~20% (Figure 5A). MukB ran as a dimer with an observed mass of 405 kDa (predicted, 346 kDa; dark grey dot). A mixture of MukB and MukF gave a major complex of 555 kDa, corresponding to a complex formed by the interaction of a MukB dimer and a MukF dimer (predicted mass, 452 kDa; blue dot). A minor faster running broad peak is likely to be a mixture of MukF dimers bound to two MukB dimers (predicted mass 797 kDa) plus ‘daisy-chain’ higher forms in which two or more MukF dimers have joined two or more MukB dimers (estimated mass towards the front of the peak 1252 kDa, corresponding to two MukF dimers bound to three MukB dimers; blue triangle).

**Figure 5.**
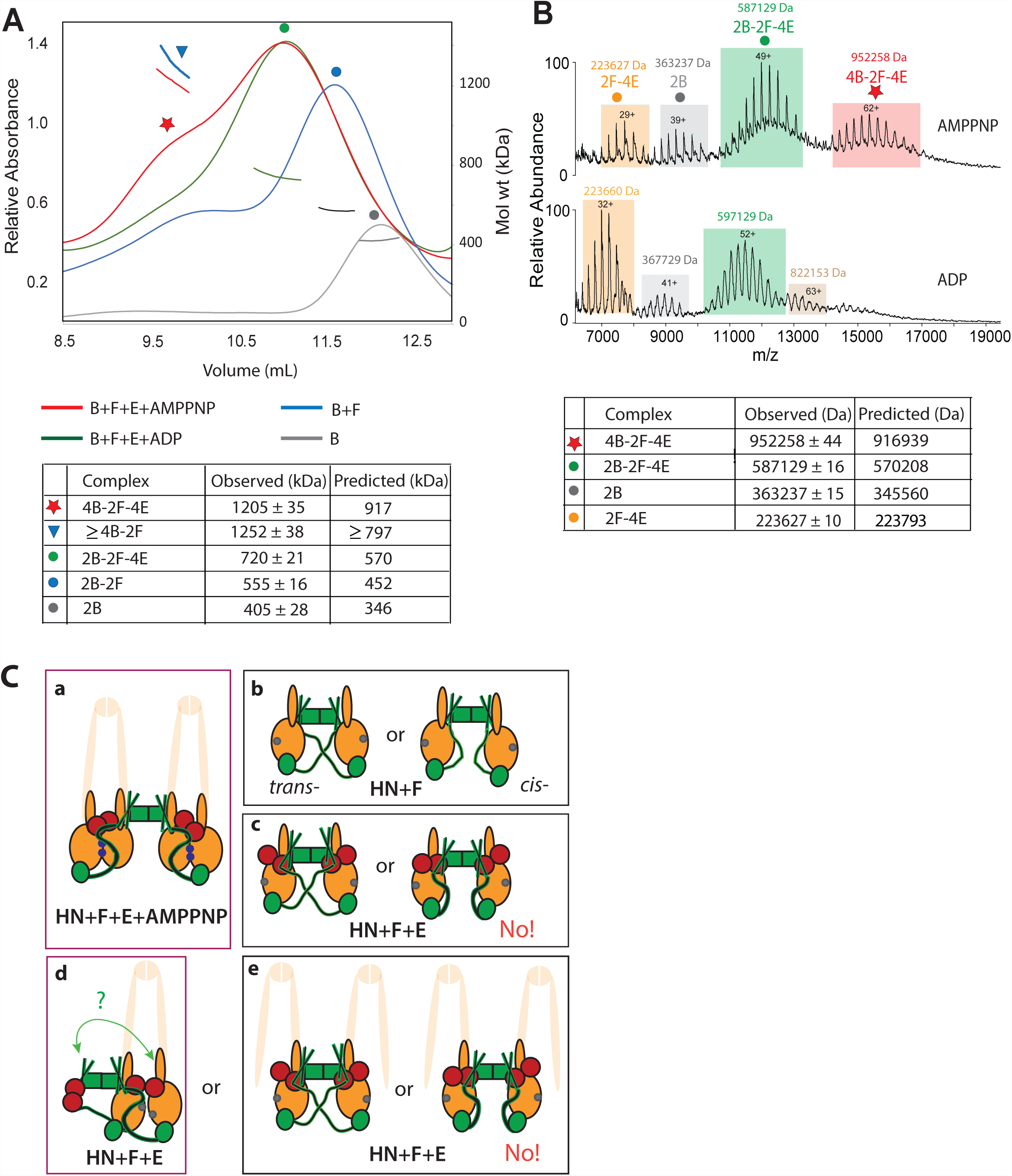
Dimers of dimers are formed with full length MukB dimers, MukEF and AMPPNP. (**A**) SEC-MALS of MukB, MukF and MukE complexes. The proteins were at concentrations 5μM B, 2.5μM F and 5μM E. Separation was through Superose 6 column. (**B**) Native mass spectra of complexes formed with MukBEF (protein concentrations as in (**A**)) in the presence of AMPPNP (top), or ADP (bottom). The predicted and observed masses are tabulated below the graphs. In (**A**), the observed masses are higher than those predicted. In the sample of B+F, this is partly because of likely higher order complexes containing additional F dimers, in addition to 4B-2F complexes (see main text). In (**B**), note that the larger species have an observed mass ~3% greater than the theoretical mass and that in the ADP sample there is a small population of observed mass 822,153 (beige); this mass is closest to a small proportion of 4B-2F or 4B-2F-E2 complexes. **(C)** Schematics of the architectures demonstrated or inferred from the biochemical analyses. The extrapolation from HN to full length MukB dimers is cartooned by showing the remainder of MukB as semi-transparent. Panel **a**, MukBEF-AMPPNP- and head engagement-dependent dimers of dimers. **b**, alternative possible architectures of HN + MukF. **c**, as **d** in the presence of MukE. The data provide evidence for the *trans*-configuration shown on the left. Panels **d** and **e**, possible configuration of 2HN-2F-4E dimers in the presence of ADP/absence of head engagement. Note that the architectures in **d** and **e**-left are topologically identical if both necks engage with a 4-helix bundle (green arrow; see Figure 4D), although if part of a full-length MukB dimer as indicated, would be a dimeric MukBEF complex (**d**) or dimer of dimer complex (**e**). ATP-bound ATPase active sites denoted as blue dots on the heads and ADP-bound or nucleotide-unbound as grey dots.

When MukB was mixed with MukF, MukE and ADP, a complex of 720 kDa was observed, consistent with the expected 2MukB-2MukF-4MukE complex (predicted mass; 570 kDa; green dot). When the same proteins were incubated with AMPPNP, along with the above peak, a smaller but substantial faster running peak of estimated mass 1205 ± 35 kDa was observed (red star). We propose that this is a 4MukB-2MukF-4MukE complex, corresponding to MukE- and AMPPNP-dependent dimers of heads-engaged dimers (predicted mass 917 kDa).

Finally, we used native mass spectrometry to analyse comparable complexes (Figure 5B). MukB, MukF and MukE were mixed and incubated with either ADP or AMPPNP. The resulting mass spectra revealed three common charge state distributions corresponding to MukB dimers (dark blue dot), MukF dimers bound by two MukE dimers (orange dot) and MukB dimers complexed with MukF dimers bound by two MukE dimers were observed (green dot). Additionally, in the sample with AMPPNP, we found a charge state series for a higher mass species that corresponds to 4MukB-2MukF-4MukE –the proposed dimer of heads-engaged MukBEF dimers (red star). Note that in the sample with ADP, there was also a small population of complexes (beige) that had a mass (822,153) most consistent with a 4MukB-2MukF complex (797 kDa).

## Discussion

Analysis of the interactions of a truncated MukB derivative (HN), containing only the ATPase head and 30% of the adjacent coiled-coil, has demonstrated that MukF dimers direct the formation of dimers of dimers dependent on AMPPNP/ATP, MukE, and MukB head engagement (Figure 5C, panel **a**). The role of MukE in part seems to be to organize and stabilize the complexes into a conformationally more compact form that allows head engagement upon ATP binding. Furthermore, MukE binding to MukF may ensure that the 4-helix bundle and C-terminal domain of a given MukF monomer bind to separate MukB heads; this *trans*-configuration thereby helping direct the formation of a kleisin-SMC tripartite proteinaceous ring (Figure 1). With the HN substrates, in the absence of engaged heads, 2HN-2F-4E dimers were the predominant form, but we cannot ascertain whether these correspond to MukBEF dimers, or dimers of MukBEF dimers (or both) (Figure 5C; compare panels **d** and **e**). Nevertheless, the analysis of full length dimeric MukB in complexes with MukFE and AMPPNP or ADP by SEC-MALS and native mass spectrometry, demonstrated that dimers of dimers form in the presence of MukBEF and AMPPNP, and that MukBEF dimers appear to be the major component in the presence of ADP, with some higher order complexes apparently forming more readily in the absence of MukE (Figure 5B).

How does the demonstration, using HN as a model for MukB, of a switch between the dimer of dimers state and the dimeric state by replacing ATP/AMPPNP by ADP relate to the *in vivo* behavior of MukBEF complexes (Figure 5C, compare panel **a** with **d**)? Because the experiments reported here use endpoint measurements in which the MukB molecules used are saturated with the added nucleotide (other than when added ATP can be hydrolysed), our assays do not assess the possible *in vivo* situation in which a dimer of dimers could undergo hydrolysis steps in both dimers simultaneously, likely leading to a MukBEF dimer (Figure 1, panel **c**, left). Alternatively, if ATP hydrolysis occurred in only one of the dimers at a given time, a dimer of dimer stoichiometry could be retained (Figure 1, panel **d**, right). Future work needs to address if both of the ATPase active sites in a MukB dimer undergo catalysis synchronously, and if there is any coordination of ATPase activity between the two MukB dimers in a dimer of dimers.

Although all four interaction interfaces between MukB and MukF (MukB neck-MukF 4HB, and MukB cap-MukF C terminal domain) can be occupied, our analyses revealed that in the absence of MukE, only two HN molecules could be bound stably at any one time. Similarly, in the presence of MukE and absence of head engagement only two HN molecules were stably bound, presumably in one of the configurations shown in Figure 4D and 5C. At present, we cannot distinguish the alternative models that can explain why only one pair of heads-engaged HN molecules can bind to a MukFE dimer. A model of MukF 4-helix bundles bound by MukB (Zawadzka et al., 2018), based on available structural information, including a ‘symmetrical juxtaposed heads’ complex with two bound MukF C-terminal domains (Woo et al., 2009), indicates that a structural constraint could prevent the interactions shown by green and blue arrows in Figure 4D, panel **a**, and the equivalent interactions in full length MukF dimers. Nevertheless, it is plausible that the modelling is misleading and interactions between both HN necks and the two 4-helix bundles do occur (Figure 5C, panel **d**, green arrows and panel **e**, left). Note that if the two necks in Figure 5C, panel **e**, left were part of the same MukB dimer, then this architecture is essentially the same as shown in panel **d**, if the interaction indicated by the green arrow occurs.

The demonstration that MukF dimers can direct the formation of dimers of heads-engaged MukB dimers in the presence of MukE and ATP/AMPPNP, using both truncated MukB derivatives and the wild-type MukB, provides strong biochemical support for our inference of such complexes in active MukBEF clusters associated with *E. coli* chromosomes *in vivo*, using quantitative imaging (Badrinaryananan et al., 2012). It therefore seems likely that all those bacteria whose genomes encode MukBEF, rather than the typical and more widely distributed SMC-ScpAB, will use a dimeric MukF to direct the formation of dimers of MukBEF dimers. Following our observation of putative dimers of dimers *in vivo*, we proposed that such complexes could be important in the transport of MukBEF with respect to chromosomal DNA by using a ‘rock-climber’ mechanism, in which the increased number of DNA-protein contact points in a dimer of dimers facilitates the transport (Badrinaryananan et al., 2012). As yet we have not succeeded in obtaining direct evidence for the putative transport mechanism. We also consider two other possibilities that are not necessarily exclusive to a role in DNA transport. First, that the role of dimer of dimer complexes is related to interaction of MukBEF with MatP-*matS* or with topoisomerase IV and the consequent biological outcomes (Nolivos et al., 2016; Zawadzki et al., 2015). Second, that dimer of dimer complexes are important for the proposed locked-phase Turing patterning mechanism that places MukBEF clusters at mid-cell or the cell quarter positions and thereby correctly positions replication origins, thereby facilitating chromosome segregation (Murray and Sourjik, 2017; Hoffman et al., 2018). It seems possible that all MukBEF orthologs use such a patterning system, along with acting in DNA transport, and hence the restriction of kleisin dimerization to MukBEF orthologs may relate to some specific property of these orthologs, other than (or in addition to) the DNA transport mechanism itself.

## Materials and Methods

### Protein purification

The proteins were expressed and purified as described (Zawadzka et al., 2018) except MgCl_2_ (1mM) was present throughout the purifications. Mutagenesis informed by the MukF dimer structure (Fennell-Fezzie et al., 2005) was used to construct a monomeric *mukF* variant.

### Size exclusion chromatography and Multi-Angle light scattering (SEC-MALS)

Proteins were mixed at respective ratios and equilibrated in 50 mM HEPES pH7.5, 100 mM NaCl, 1 mM MgCl_2_, 1 mM DTT and 10% glycerol buffer supplemented with 1 mM ADP, ATP, or AMPPNP for 3 hours in. 100 µL of these mixtures were loaded onto either a Superose 6 HR10/30 column, or a Superdex 200 HR10/30 column (GE), equilibrated with the same buffer lacking glycerol and nucleotides. The separation was conducted at flow rate of 0.5 mL/min. Presence of DTT or TCEP as reductants did not influence the results. SEC-MALS analysis was performed at 22 °C using a Shimadzu (Kyoto, Japan) chromatography system, connected in-line to a Heleos8+ multi angle light scattering detector and an Optilab T-rEX refractive index (RI) detector (Wyatt Technologies, Goleta, CA). Results were processed and analysed using ASTRA 6 (Wyatt Technologies).

### Native and SDS polyacrylamide gel electrophoresis (PAGE)

SDS polyacrylamide gels were prepared as described (Zawadzka et al., 2018). 6% native polyacrylamide gels were poured in 125 mM Tris buffer pH 8.8. Gels were run using Tris-Glycine running buffer (25 mM Tris-Cl, 192 mM glycine). Purified proteins were mixed at respective ratios in 50 mM HEPES pH 7.5, 100 mM NaCl, 1mM MgCl_2_, 1 mM DTT along with respective nucleotide (1 mM ADP/ATP or AMPPNP) and equilibrated for 3 hours at room temperature. Samples were mixed with 20% glycerol before loading. Gels were run at 35 mA for 30-35 minutes and stained using Instant Blue.

### Isothermal calorimetry (ITC)

Reaction samples containing MukE (400 µM) and MukF (20 µM) were equilibrated in 50 mM HEPES pH 7.5 100 mM NaCl, 1mM MgCl_2_ and 1 mM DTT. Binding was assayed in a Malvern PEAQ ITC instrument at 25°C. Averages and standard deviations of the obtained parameters are reported from triplicate experiments. Data were analysed using the manufacturer’s software assuming a single binding site model.

### Native-state ESI-MS spectrometry

Prior to MS analysis, protein samples were buffer-exchanged into 200 mM ammonium acetate pH 8.0, using a Biospin-6 (BioRad) column and introduced directly into the mass spectrometer using gold-coated capillary needles (prepared in-house) [DOI: 10.1038/nprot.2007.73]. Data were collected on a modified QExactive hybrid quadrupole-Orbitrap mass spectrometer (Thermo-Fisher Scientific) optimized for analysis of high-mass complexes, using methods previously described [DOI: 10.1038/nmeth.3771]. The instrument parameters were as follows: capillary voltage 1.2 kV, S-lens RF 100%, quadrupole selection from 2,000 to 20,000 m/z range, collisional activation in the HCD cell 200 V, argon UHV pressure 1.12 × 10−9 mbar, temperature 60 °C, resolution of the instrument 17,500 at m/z = 200 (a transient time of 64 ms) and ion transfer optics (injection flatapole, inter-flatapole lens, bent flatapole, transfer multipole: 8, 7, 6 and 4 V, respectively). The noise level was set at 3 rather than the default value of 4.64. No in-source dissociation was applied. Where required, baseline subtraction was performed to achieve a better-quality mass spectrum.

### Contributions

The project was conceived and directed by DJS, LKA and KR. Experiments were undertaken by KR and MT. RB, KZ, FW and OK prepared key reagents. JRB and CVR performed mass spectrometry analysis. The manuscript was written by KR, LKA and DJS.

## Acknowledgements

We thank the Biochemistry Biophysical facility and its manager, Dr. D. Staunton, for providing access to ITC and assistance with SEC-MALS. The research was funded by a Wellcome Investigator Award to DJS (200782/Z/16/Z). Research in the CVR group is supported by an MRC Programme grant (MR/N020413/1), an ERC Advanced Grant ENABLE (695511) and a Wellcome Investigator Award (104633/Z/14/Z).

---

**Figure 3, supplementary figure 1.**
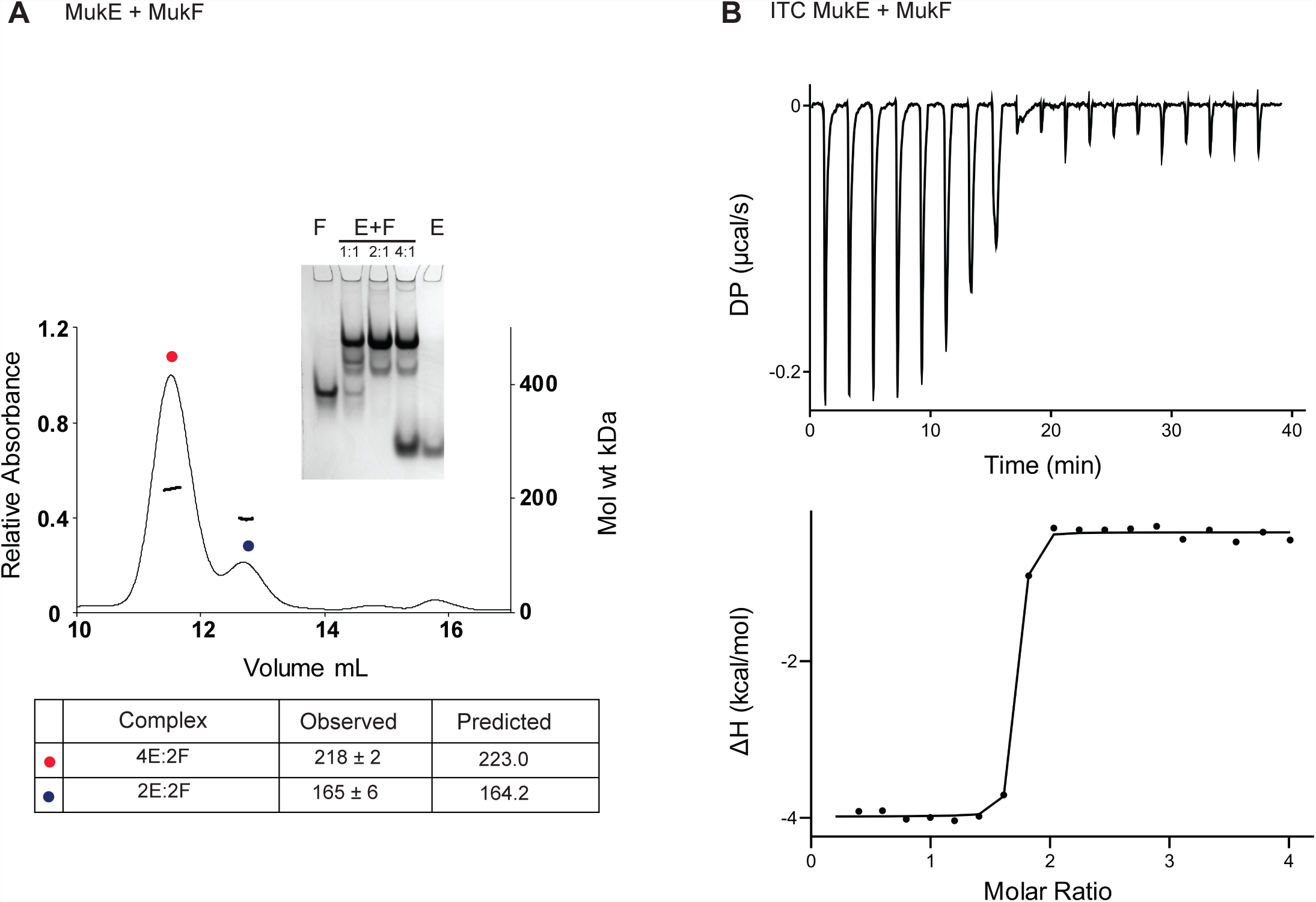
**A.** SEC-MALS of MukE-MukF complexes on Superdex 200 column with 100 µL of 10 µM MukF and 20 µM MukE. Inset figure shows a 6 % Native gel electrophoresis. Lanes from left to right are 5 µM MukF, 5 µM MukE + 5 µM MukF, 10 µM MukE + 5 µM MukF, 20 µM MukE + 5 µM MukF, 5 µM MukE. **B.** ITC binding isotherm of 400 µM MukE titrated into 20 µM MukF at 25 °C in an ITC 200. Fitted parameters for a single binding site model from three independent measurements were *N*=1.87 ± 0.24, *K*_d_=6.97 ± 2.6 nM, Δ*H*=−14.5 ± 0.15 kcal mol^−1^, TΔ*S*= −32.1 kJ mol^−1^ MukE monomers bound to a middle region of MukF have interacting surfaces with both monomers of a MukB head (total buried surface area of 344 and 630 Å^2^; Woo et al., 2009)

**Figure 3, supplementary figure 2.**
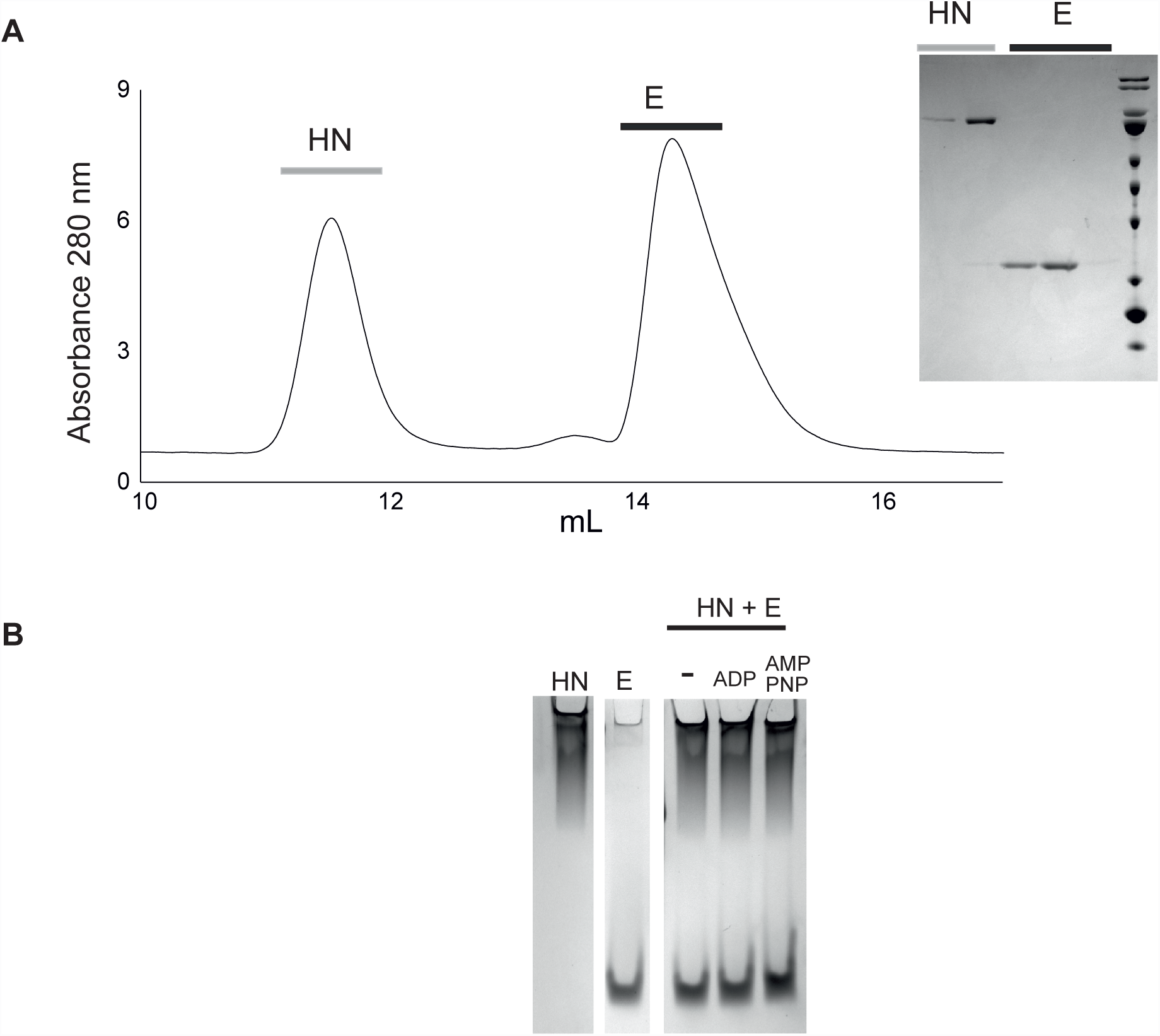
**A.** SEC of mixed HN (10 µM) and MukE (10 µM). Inset, SDS PAGE of peak fractions **B.** Native gel electrophoresis of HN + MukE, with and without the indicated nucleotides.

**Figure 4, supplementary figure 1.**
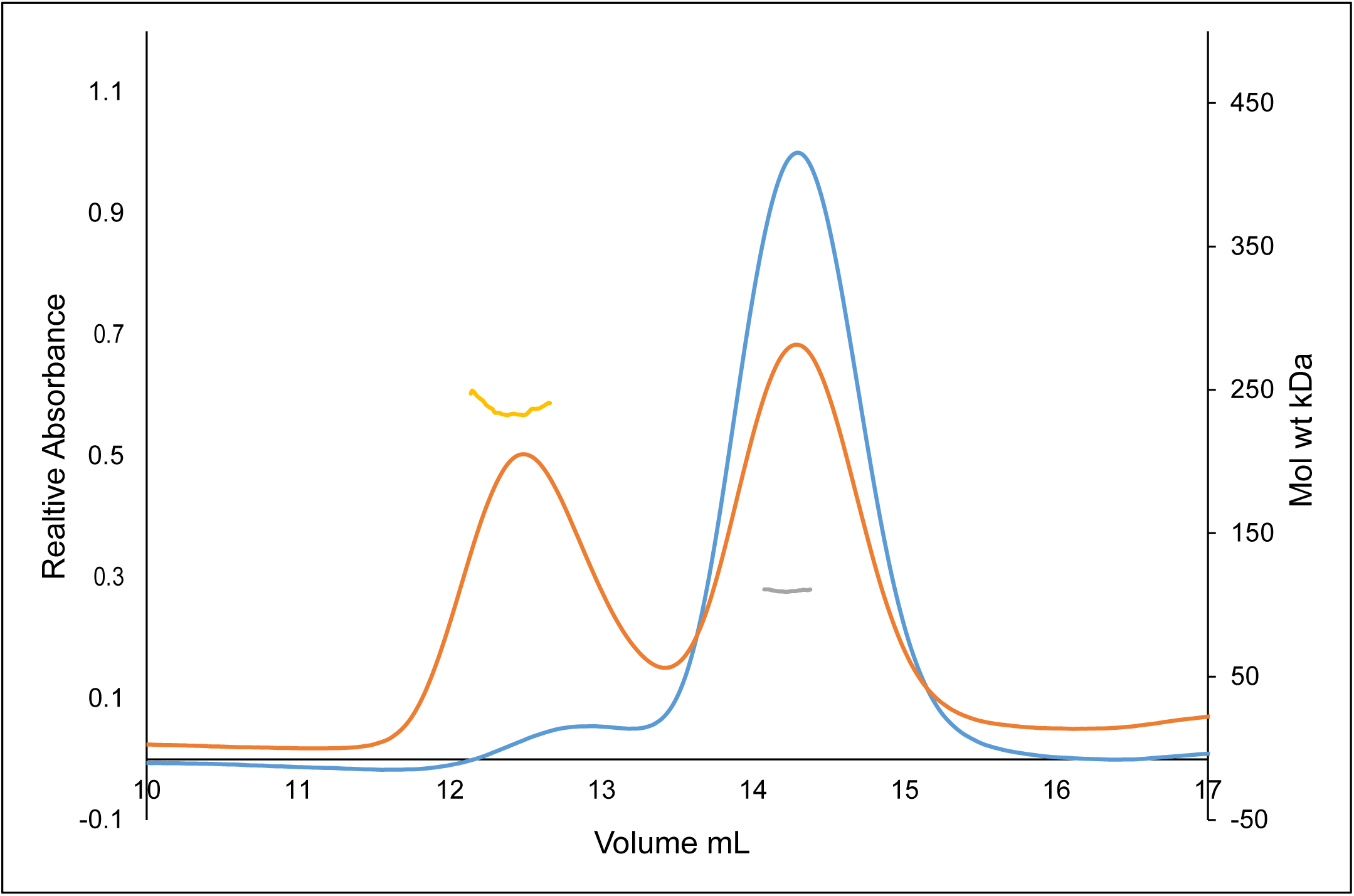
HN^EQ^ dimerises independently of MukF. Superdex 200 SEC-MALS analysis of HN^EQ^ preincubated with ATP (orange) or ADP (blue).

## References

Badrinarayanan A, Reyes-Lamothe R, Uphoff S, Leake MC, Sherratt DJ. (2012) In vivo architecture and action of bacterial structural maintenance of chromosome proteins Science 338: 528–531.

Brézellec P, Hoebeke M, Hiet MS, Pasek S, Ferat JL. (2006) DomainSieve: a protein domain-based screen that led to the identification of dam-associated genes with potential link to DNA maintenance Bioinformatics 22: 1935–1941.

Bürmann F, Shin HC, Basquin J, Soh YM, Gimenez-Oya V, Kim YG, Oh BH,Gruber S. (2013) An asymmetric SMC-kleisin bridge in prokaryotic condensin Nature Structural & Molecular Biology 20: 371–379.

Danilova O,Reyes-Lamothe R, Pinskaya M, Sherratt D, Possoz C. (2007) MukB colocalizes with the oriC regionand is required for organization of the two Escherichia coli chromosome arms into separate cell halves. Molecular Microbiology 65:1485–1492.

Fennell-Fezzie R, Gradia SD, Akey D, Berger JM. (2005) The MukF subunit of Escherichia coli condensin: architecture and functional relationship to kleisins. Embo Journal 24: 1921–1930.

Ganji M, Shaltiel IA, Bisht S, Kim E, Kalichava A, Haering CH, Dekker C. (2018) Real-time imaging of DNA loop extrusion by condensin. Science 360: 102–105.

Gligoris TG, Scheinost JC, Burmann F, Petela N, Chan KL, Uluocak P, Beckouet F, Gruber S, Nasmyth K, Lowe J. (2014) Closing the cohesin ring: structure and function of its Smc3-kleisin interface. Science 346: 963–967.

Hirano T. (2016) Condensin-based chromosome organization from bacteria to vertebrates Cell 164: 847–857.

Hofmann A, Mäkelä J, Sherratt D, Heermann D, Murray SM (2018) Self-organized segregation of bacterial chromosomal origins bioRxiv.org doi: https://doi.org/10.1101/304600.

Huis in ‘t Veld PJ, Herzog F, Ladurner R, Davidson IF, Piric S, Kreidl E, Bhaskara V, Aebersold R, Peters JM. (2014) Characterization of a DNA exit gate in the human cohesin ring. Science 346:, 968–972.

Lammens A, Schele A, Hopfner KP. (2004) Structural biochemistry of ATP-driven dimerization and DNA-stimulated activation of SMC ATPases Current Biology 14:1778–1782.

Murray SM, Sourjik V. (2017) Self-organization and positioning of bacterial protein clusters. Nature Physics 13: 1006.

Nasmyth K. (2017) How are DNAs woven into chromosome. Science 358: 589–590.

Nicolas E, Upton AL, Uphoff S, Henry O, Badrinarayanan A, Sherratt D. (2014) The SMC complex MukBEF recruits topoisomerase IV to the origin of replication region in live Escherichia coli. mBio 5, e01001–01013.

Nolivos S, Sherratt D. (2014) The bacterial chromosome: architecture and action of bacterial SMC and SMC-like complexes. FEMS Microbiology Reviews 38: 380–392.

Nolivos S, Upton AL, Badrinarayanan A, Muller J, Zawadzka K, Wiktor J, Gill A, Arciszewska L, Nicolas E, Sherratt D. (2016) MatP regulates the coordinated action of topoisomerase IV and MukBEF in chromosome segregation. Nature Communications 7: 10466.

Uhlmann F. (2016) SMC complexes: from DNA to chromosomes. Nature Reviews Molecular Cell Biology 17: 399–412.

Upton AL, Sherratt DJ. (2013) Breaking symmetry in SMCs. Nature Structural & Molecular Biology 20: 246–249.

Woo JS, Lim JH, Shin HC, Suh MK, Ku B, Lee KH, Joo K, Robinson H, Lee J, Park SY, Ha NC, Oh BH. (2009) Structural studies of a bacterial condensin complex reveal ATP-dependent disruption of intersubunit interactions Cell 136: 85–96.

Zawadzka K, Zawadzki P, Baker R, Rajasekar KV, Wagner F, Sherratt DJ, Arciszewska LK (2018) MukB ATPases are regulated independently by the N- and C-terminal domains of MukF kleisin. eLife 7.

Zawadzki P, Stracy M, Ginda K, Zawadzka K, Lesterlin C, Kapanidis AN, Sherratt DJ. (2015) The Localization and Action of Topoisomerase IV in Escherichia coli Chromosome Segregation Is Coordinated by the SMC Complex, MukBEF Cell Reports 13:2587–2596.

